# Identification of a Chimeric M4 Protein and Novel *Emm* Pattern in Currently Circulating Strains of *Emm4* Group A *Streptococcus*

**DOI:** 10.1101/333666

**Authors:** Sruti DebRoy, Xiqi Li, Awdhesh Kalia, Jessica Galloway Peña, Brittany J. Shah, Vance G. Fowler, Anthony R. Flores, Samuel A. Shelburne

## Abstract

Group A *Streptococcus* (GAS) is classified by sequence of the gene encoding the M protein (*emm*) and the patterns into which *emm* types are grouped. We discovered a novel *emm* pattern in *emm4* GAS, historically considered pattern E, arising from a fusion event between *emm* and the adjacent *enn* gene. We identified the *emm/enn* fusion event in 51/52 *emm4* GAS strains isolated by national surveillance in 2015. GAS isolates with an *emm/enn* fusion event completely replaced pattern E *emm4* strains over a 4-year span in Houston (2013-2017). The novel *emm/enn* gene fusion and new *emm* pattern has potential vaccine implications.

## INTRODUCTION

Group A *Streptococcus* (GAS) is among the most ubiquitous human pathogens and is classified into *emm* types based on the 5’ sequence of the *emm* gene that encodes the hypervariable M protein, a key GAS virulence determinant [1]. GAS infection is thought to engender serotype-specific immunity, and the M protein or its domains are leading vaccine candidates [2]. In addition to invariably containing the *emm* gene, the *emm* region can also contain one or two *emm*-like genes which are typically designated as *mrp* and *enn* (Figure 1A) [3]. The 3’ end of the *emm* and *emm*-like genes encode a peptidoglycan–spanning (PG) domain, and PG sequence variation determines four different subfamilies (SF-1 through SF-4, Figure 1A). The composition of *emm*-family genes along with their respective PG domain subfamily types divides GAS strains into five *emm* patterns (A-E), and for the overwhelming majority of GAS strains studied to date, there is strict concordance between *emm* type and *emm* pattern [3]. The *emm* pattern of GAS strains strongly correlates with the preferred epithelial site of infection i.e. pharynx versus skin [3]. Thus, *emm* pattern A-C strains are considered “throat specialists”, whereas pattern D are “skin specialists”, and pattern E are “generalists” [4].

**Figure 1.**
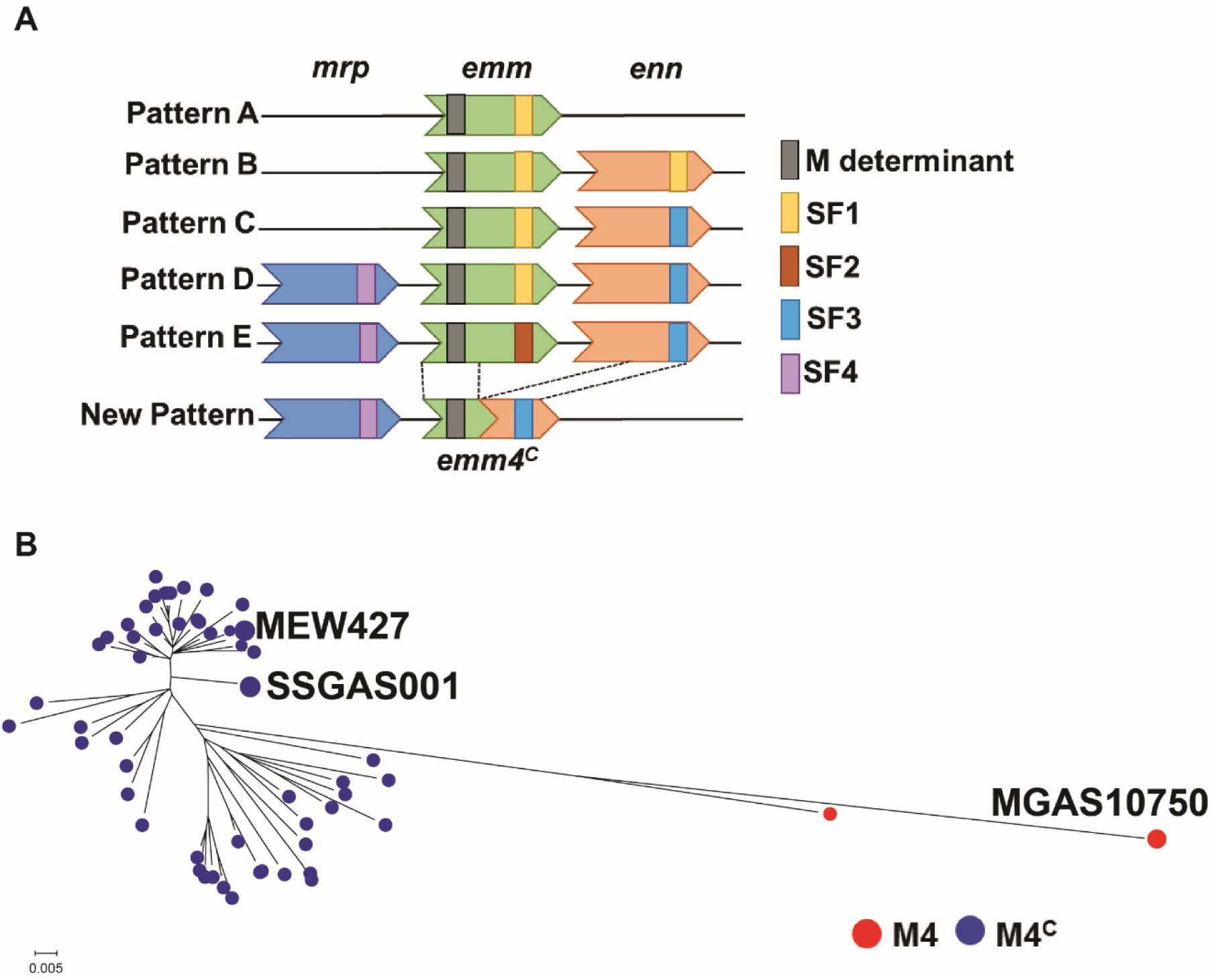
Characterization of GAS isolates. (A) Schematic diagram to compare the new *emm* pattern to existing patterns (adapted from reference [3]). The *emm* type-specific determinant and the subfamily of the PG domain is indicated. (B) Maximum-likelihood tree reconstructed on 814 core SNPs of *emm*4 GAS strains from CDC [11] along with publicly available *emm4* GAS genomes (names are provided). The type of M protein is color coded as indicated in legend.

The majority of data establishing the five *emm* patterns and their relationship to *emm* type were generated from strains isolated in the 1980s and 1990s using targeted approaches such as PCR and Southern blot [4, 5]. Initial whole genome sequencing (WGS) studies of one or a few GAS strains per *emm* type done in the 2000s have supported these findings [6]. However, the past decade has witnessed a marked increase in WGS of large cohorts of GAS strains which has revealed significant intra-*emm* type diversity [7]. Moreover, the highly antigenic and thus hypervariable nature of the M protein has long been recognized [8]. Therefore, with the additional density of sequencing data and the elapsed time since the recognition of the five *emm* patterns, one might predict that additional variations in the *emm* region would be recognized. Herein, we report the identification of genetic events in currently circulating *emm4* strains, which were previously considered pattern E, that give rise to a chimeric M protein and a novel GAS *emm* pattern.

## MATERIALS AND METHODS

The invasive GAS *emm4* strain SSGAS001 was isolated from a point-source outbreak in 2015 and its genome described using the strain name “Duke small” [9]. We accessed the complete genomes of the 2001 pharyngeal isolate MGAS10750, considered the *emm4* reference strains, and the 2015 pharyngeal *emm4* isolate MEW427 from NCBI [6, 10]. Short-read sequencing data of invasive *emm4* strains collected by the Centers for Disease Control and Prevention (CDC) during 2015 [11] were accessed from BioProject PRJNA395240 on the short-read archive. *Emm* region sequences were extracted from annotated draft genomes and visualized through Geneious (Biomatters Ltd, NJ, USA). Two core single nucleotide polymorphism (SNP)-based methods were deployed to reconstruct WGS-based phylogenies: a) kSNP v3.0 identified core SNPs based on reference free k-mer analysis (k-mer size of 19, default settings). These SNPs were used to reconstruct a maximum parsimony tree; and b) parsnp v1.2 aligned genomes to the reference, identified SNPs in local collinear blocks, and subsequently reconstructed an approximate maximum-likelihood tree using FastTree 2. Due to similar findings, only trees reconstructed by parsnp are provided. 88 *emm*4 GAS were identified among a total of 930 sterile and non-sterile GAS isolates collected at Texas Children’s Hospital in Houston, TX between 2013 and 2017 under a protocol approved by the Institutional Review Board at Baylor College of Medicine. The *emm* region was amplified by PCR and subject to Sanger sequencing (primer sequences are in Supplemental Table 1) with subsequent identification of the *emm/enn* gene fusion.

## RESULTS

*Emm4* strains commonly cause both invasive and non-invasive GAS disease and are considered a prototype *emm* pattern E GAS [1, 12]. We previously compared the complete genome of an invasive *emm4* strain isolated in 2015 (strain SSGAS001) to that of the reference *emm4* strain MGAS10750, which was isolated in 2001 [6, 9]. Closer examination revealed a fusion event between *emm* and the immediately downstream *enn* gene in SSGAS001 resulting in a chimeric *emm* gene, which we have designated as *emm4^C^*. The *emm4^C^* gene is a fusion of the 5’ end of the *emm* and the 3’ end of the *enn* gene of MGAS10750 (Figure 1A). Given that the 5’ of *emm4^C^* is unchanged, SSGAS001 is still categorized as *emm4*. However, the C-terminal half of the *emm4^C^*-encoded protein, M4^C^, only shares 62% identity with the C-terminus of the MGAS10750 M4 protein. Moreover, the chimeric M4^C^ protein now contains an SF-3 allele in the PG domain rather than the SF-2 allele found in the PG domain of the MGAS10750 M protein. In addition to generating a new chimeric M protein, the *emm/enn* gene fusion event observed in SSGAS001 also results in just two genes in the *emm* region (*mrp* and *emm*) instead of the three (*mrp, emm* and *enn*) that are characteristic of Pattern E GAS (Figure 1A). Thus, the gene fusion event gives rise to an *emm* pattern heretofore undescribed in GAS strains. We analyzed the other fully sequenced *emm4* GAS strain, MEW427, also isolated in 2015 [10], and found it to have the identical *emm4^C^* gene and *emm* pattern as strain SSGAS001.

Next, we wanted to determine the frequency of the gene fusion event amongst currently circulating *emm*4 GAS strains. We first analyzed published WGS data from 52 invasive *emm4* isolates collected by the CDC in 2015 [11]. We identified the *emm*/*enn* gene fusion event in 51/52 strains with only a single strain containing *emm* pattern E (Figure 1B, Supplemental Table 2).

Whole genome-based phylogenomics revealed that strains with the *emm/enn* gene fusion clustered distinctly from strains with the canonical *emm4* gene (Figure 1B). We identified six variations of the M4^C^ protein (Figure 2A, Supplemental Figure 1). The 350 amino acid long variant, which we have named M4^C-350^, occurred most commonly and was identical to M4^C^ in strains SSGAS001 and MEW427. M4^C-350^ differs from the other common M4^C^ variant, M4^C-392^, by a 126 nucleotide (42 amino acid) deletion (Figure 2A). Together M4^C-350^ and M4^C-392^ are present in 90% of strains in the CDC dataset that have the *emm*/*enn* gene fusion event (Figure 2B). The other four variants appear to be derived via insertions or deletions in the M4^C-392^ or M4^C-350^ protein (Supplemental Figure 1).

**Figure 2.**
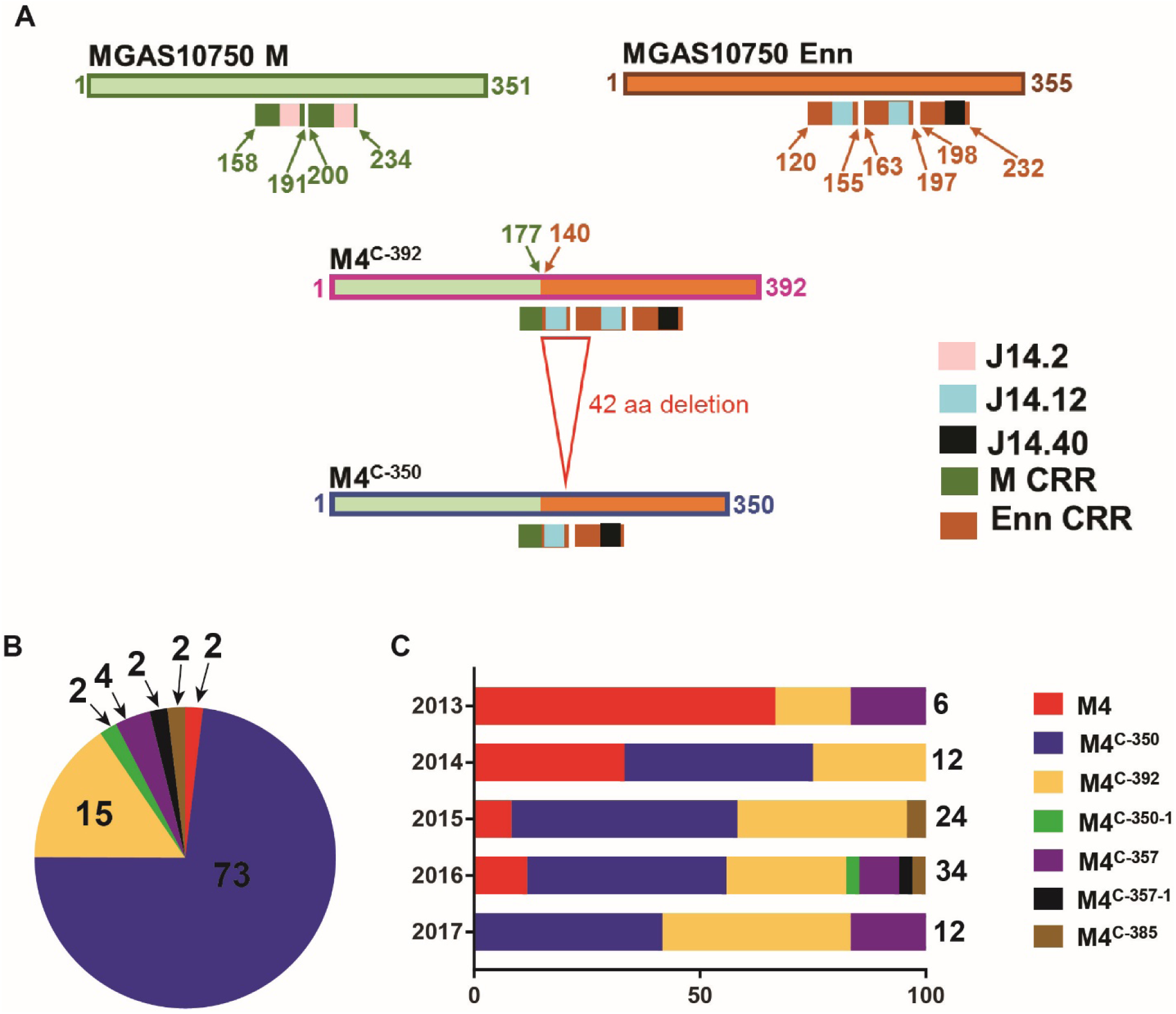
(A) Schematic diagram showing the two most common chimeric M protein variants. The fusion point of the *emm* and *enn* genes is indicated by the change in box shading with numbers indicating the last amino acid from *emm* and first amino acid from *enn* respectively in M4^C^ protein. The 42 amino acid (126 nucleotide) deletion that is the difference between M4^C-392^ and M4^C-350^ is noted. C-terminal repeat regions (CRRs) along with J14 sequences are displayed as color-coded boxes. (B) Percent distribution of the canonical and chimeric M4 proteins in 52 *emm*4 GAS strains collected by the CDC in the United States in 2015 depicted as a pie-chart. Numbers indicate % of total. (C) Distribution of the canonical M4 and the different M4^C^ variants identified in 88 *emm4* strains isolated from patients at Texas Children’s Hospital between 2013 and 2017 expressed as a percentage of the number of isolates per year. Total number of isolates in a given year is indicated on the right of the bars. There are relatively fewer isolates in 2013 because collection only began in May, 2013. For (B) and (C), the type of M protein is color coded as indicated in legend.

Although the N-terminal of the M4^C^ proteins are identical to the canonical M4 protein, the M4^C^ proteins have different C-terminal repeat regions (CRRs). The CRRs represent a domain within the cell surface-exposed portion of the M and Enn proteins that contain tandemly arranged blocks of direct sequence repeats which have been targeted as potential GAS vaccine candidates [13]. The sequence repeats are not always identical and can be further classified based on the sequence of the C-terminal 14 amino acids (J14) in a given repeat [14]. The MGAS10750 M4 protein contains two CRRs, both of which have the J14.2 sequence. Unlike MGAS10750, M4^C-392^ contains three CRRs, but all other M4^C^ variants also possess two CRRs each. However, the composition of the CRRs in every M4^C^ variant differs from that in MGAS10750 in terms of the J14 sequence (Figure 2A, Supplemental Table 2, Supplemental Figure 1). Importantly, the M4^C^ proteins are present in strains isolated from multiple locations in the United States rather than being confined to a single location (Supplemental Figure 2).

To begin to assess temporal emergence of *emm4* strains with the chimeric M protein, we analyzed 88 *emm4* isolates collected at the Texas Children’s Hospital in Houston between 2013 and 2017 via Sanger sequencing of the *emm* region (Supplemental Table 3). We identified the *emm/enn* gene fusion in 74 strains (84%). While 66% of the *emm4* strains isolated in 2013 were still the canonical *emm* pattern E, there was total replacement by strains carrying *emm4^C^* by 2017 (Figure 2C). Similar to the CDC strains, the M4^C-350^ and M4^C-392^ variants were most common among the Houston isolates. Consistent with the hypothesis that M4^C-350^ arose from M4^C-392^, M4^C- 350^-containing strains were not detected until one year after the initial identification of strains with the M4^C-392^ protein (Figure 2C).

## DISCUSSION

Since its discovery some 90 years ago, M protein has been considered the key GAS virulence determinant and is currently a prime vaccine candidate [2]. Despite extensive investigations over the past decades, continued study of M protein and the *emm* region continues to uncover new findings. Herein, we demonstrate that >90% of recently circulating *emm4* GAS strains contain a non-canonical chimeric M4 protein and a previously unrecognized *emm* pattern.

The key finding of this study was identification of a gene fusion event that gave rise to three novel entities. First, it created a chimeric *emm* gene, *emm4^C^*, that consists of the 5’ end of the reference *emm4* gene and the 3’ end of the *enn4* gene. Second, formation of the chimeric *emm4^C^* gene eliminates the *enn* gene thereby producing an *emm* region that only contains *mrp* and *emm*. Finally, unlike previously reported M proteins which harbor either the SF-1 or SF-2 allele in the PG domain [3], M4^C^ carries the SF-3 allele of the PG subfamily. To our knowledge, our study is the first report of a fusion event between *emm* family genes that alters the M protein as well as the first record of an *emm* pattern different from the five established ones.

Analysis of 140 recent *emm*4 isolates revealed that strains with the chimeric *emm4^C^* gene and the new *emm* pattern have almost completely replaced the previously circulating pattern E strains. This finding implies that the gene fusion event confers an advantage to the strains with the novel *emm* pattern or that strains with the gene fusion event have additional changes driving their proliferation. Given the critical nature of M protein in GAS host-pathogen interaction, it is plausible that the gene fusion event is at least a partial catalyst for the observed replacement. Multiple key aspects of GAS pathogenesis have been ascribed to M protein, such as binding of host molecules and inhibiting phagocytosis, although the precise function differs depending on M protein composition [1]. Additionally, M protein is the dominant GAS target of the human immune system [2]. Thus, the theoretical advantage conferred by M4^C^ protein could be due to augmented function of the chimeric M protein and/or to an improved ability to evade the host immune system.

In terms of altered susceptibility to human immunity, M protein sequence variation can confer changes in immune recognition and has been associated with clonal population expansion [7]. However, reports of M protein variation have described small-scale changes (i.e. insertions, deletions, tandem repeat variation) in the N-terminal region whereas we observed replacement of the entire C-terminus [7, 8]. Although the N-terminus is typically considered to be the portion of the M protein under selective immune pressure [1], there are also data supporting immunogenicity of the C-terminal region. Specifically, J14 sequences in the CRRs have been shown to induce immunity to GAS [13], and the *emm*/*enn* fusion events alter the J14 sequence in every variant of the M4^C^ protein that we observed (Figure 2A). The *enn* gene has been reported to be transcribed at very low levels and may not be translated at all due to mutations early in the coding region [15]. Therefore, we speculate that previously circulating *emm4* strains bearing the classic pattern E and canonical *emm* may not have engendered an immune response to the portion of the Enn protein now present in the M4^C^ protein. Hence, these data suggest that the *emm/enn* gene fusion may have significant immunological effects on GAS-host interactions, but this needs to be experimentally tested.

In summary, we report the occurrence of a recombination event that has given rise to a chimeric *emm* gene and a novel *emm* pattern and provide evidence of its predominance among current *emm4* strains. These data suggest that GAS mechanisms that alter the M protein are more varied than previously appreciated which could impact the efficacy of M protein-based vaccine strategies.

## Acknowledgements

We thank Dr. Bernie Beall for critical reading of the manuscript and suggestions regarding the origin of the different M protein variants.

